# Orb-web spider color polymorphism through the eyes of multiple prey and predators

**DOI:** 10.1101/160341

**Authors:** Nathalia Ximenes, Felipe M. Gawryszewski

## Abstract

The sensory drive theory predicts that signals, sensory systems, and signaling behavior should coevolve. Variation in the sensory systems of prey and predators may explain the diversity of color signals, such as color polymorphism. The spider *Gasteracantha cancriformis* (Araneidae) possesses several conspicuous color morphs. The aim of the present study was to assess whether the color polymorphism of *G. cancriformis* may be maintained by pressure from multiple signal receivers, such as prey and predators with distinct color vision systems. Although, the multiple receivers world is a more realistic scenario, it has received little attention. In orb-web spiders, the prey attraction hypothesis states that conspicuous colors are prey lures that increase spider foraging success via flower mimicry. However, in highly defended species, conspicuous colors could also be a warning signal to predators. We used color vision modelling to estimate chromatic and achromatic contrast of *G. cancriformis* morphs as perceived by potential prey and predator taxa. Our results revealed that individual prey and predator taxa perceive the conspicuousness of morphs differently. For instance, the red morph is perceived as quite conspicuous to lepidopteran prey and avian predators, but not by other insects. Therefore, the multiple prey and predator hypotheses may explain the evolution of color polymorphism in *G. cancriformis*. However, flower mimicry hypothesis was weakly corroborated. Other parameters that are not evaluated by color vision models, such as distance, shape, angle, and pattern geometry could also affect the perception of color morphs by both prey and predators and thereby influence morph survival.

## INTRODUCTION

The evolution and maintenance of color polymorphism have traditionally been attributed to apostatic selection (Clarke, 1979). Assuming that predators form a search image (Tinbergen, 1960), the advantage of rarity promotes the coexistence of multiple prey types and stabilizes polymorphisms (Bond 2007). Nonetheless, other adaptive and non-adaptive explanations for the evolution and maintenance of color polymorphisms have been proposed (Gray and McKinnon, 2007). For instance, gene flow between populations with distinct phenotypes that are favored by natural selection could maintain polymorphism within populations (Farkas et al., 2013; Gray and McKinnon, 2007).

In the context of visual signaling, the distinct visual systems of prey and predators may play a role in the evolution and maintenance of color polymorphisms (Ruxton et al., 2004; White and Kemp, 2015). Animal communication involves the generation, emission transmission, and processing of the signal by a receiver, in which an appropriate response is elicited (Endler 1993). Any factors that affect these steps can influence signal efficiency and, as a result, affect the direction of communication evolution (Endler 1993). Thus, the diversity of signals are likely influenced by variation in the sensory systems of receivers.

Many orb-web spiders exhibit conspicuous coloration. Although sexual selection is a common explanation for bright coloration in other taxa such as birds (Ryan, 1990), this scenario is less likely to happen in orb web spiders, because they have limited visual acuity (Foelix, 2011). *Argiope argentata* (Araneidae), for instance, seems to possess only one photoreceptor (Tiedemann, 1986). The prey attraction hypothesis states that the bright coloration of some spiders lures insects, possibly by mimicking flower coloration (e.g. Craig and Ebert, 1994; Hauber, 2002). The hypothesis has been empirically tested several times, and most studies have found support for it. The polymorphic *Nephila pilipes* (Nephilidae) present a melanic and a bright colored morph (Tso et al., 2004). The bright color patterns of this species are thought to resemble symmetric flower patterns that may attract bees, owing to the innate preference of bees for symmetry (Chiao et al., 2009). Moreover, yellow patches on the spider’s body may be perceived as food resources by flowers visitors (Tso et al. 2004). Besides being attractive to pollinators, the yellow patches on the species’ body also seems to attract hymenopteran predators. Therefore, it is possible that there is a trade-off between foraging success and predation risk in polymorphic populations in which some morphs are more cryptic than others (Fan et al., 2009).

The predators of orb-web spiders possess very distinct visual systems. Birds, for example, are tetrachromats, whose photoreceptors are most sensitive to ultraviolet-violet, blue, green, and red (Hart 2001), whereas spider hunting wasps, such as members of the Sphecidae, are trichromats, whose photoreceptors are most sensitive to ultraviolet, blue, and green (Peitsch, 1992; Briscoe and Chittka, 2001). Similarly, the insect prey of orb-web spiders also vary in their types of color vision. For example, bees are trichromats with spectral sensitivities that are similar to those of sphecid wasps (Briscoe and Chittka, 2001), whereas some lepidopterans are tetrachromats, and some dipterans possess photoreceptors with five different sensitivity peaks (Schnaitmann et al., 2013). Therefore, the maintenance of spider color polymorphism may result not only from a trade-off between prey attraction and capture success but also from selective pressure from multiple receivers (Endler, 1992; Ruxton et al., 2004; White and Kemp, 2015)

The orb-web spider *G. cancriformis* constructs large webs and rests in the web hub during the day (Levi, 1978). Females of the species possess a hard abdomen with three pairs of spines and vary in color, with some morphs quite conspicuous to human observers (Levi, 1978; Gawryszewski and Motta, 2012). The ventral side of females are mostly black, sometimes with small bright spots. In one studied population, the dorsal side of females possessed black or reddish spines and four different color patterns: yellow, white (without UV reflectance), red, and a combination of black and white (white patches reflects UV; Gawryszewski 2007; Gawryszewski and Motta, 2012). Adult females measure from 5 to 7 mm in length and 10 to 13 mm in width (Muma, 1971), whereas the males are brownish, small, and do not exhibit chromatic variation (Levi, 1978). The prey attraction hypothesis does not seem to explain the coloration of the orb-web spider *Gasteracantha cancriformis* (Araneidae), since both naturally bright morphs and yellow-painted individuals failed to capture more prey than either naturally cryptic morphs or black-painted individuals (Gawryszewski and Motta, 2012). Nonetheless, it remains the possibility that each color morphs attracts preferentially specific types of prey. Furthermore, although evidence is still needed, Edmunds and Edmunds (1983) suggested that the conspicuous body coloration of *Gasteracantha* spiders might serve as a warning signal to predators.

Considering that the same “color” may be perceived as cryptic or conspicuous by different species (Endler and Mappes 2004), each color morph of a polymorphic populations may represent an adaptation to particular visual systems of prey or predator species (Endler, 1992; Ruxton et al., 2004; White and Kemp, 2015). Therefore, it is plausible that the variation of color among individuals within a population is affected by a diverse range of interactions that leads to different selection process. To date, the role of multiple predators on the evolution of prey coloration has been approached by theoretical models (Endler and Greenwood, 1988; Endler and Mappes, 2004). Endler and Greenwood (1988) models, for instance, indicated that a stable polymorphisms might evolve in the presence of anti-apostatic (positive frequency-dependent) from different predators, given that predators perceive prey conspicuousness differently and there is a covariance between the relative degree of crypsis and the degree of frequency-dependent selection by each predator.

In this study we aimed to explore old and new hypothesis that could potentially explain the maintenance of color polymorphism in a spider species. The aim of the present study was to investigate three hypotheses for the evolution and maintenance of color polymorphism, using *G. cancriformis* as a model. Two derivations from the prey attraction hypothesis include (1) the *multiple prey hypothesis*, which posits that each color morph is adapted to lure a specific type of prey, which posits that the spiders attract prey *via* aggressive mimicry of flower colors and that each color morph mimics a different flower color. In addition, (3) the *multiple predator hypothesis* posits that the conspicuous colors found in spiders could serve as warning signals to predators and that color polymorphism could evolve and be maintained if each color morph is adapted to the vision of a specific predator.

## MATERIALS AND METHODS

### Color vision model

Color perception depends on both the signal reflectance and observer visual system, as well as on the background reflectance spectrum and ambient light intensity (Endler 1990). Physiological models of color vision include all these factors and have been effective for objectively studying animal coloration (i.e., independent of human subjective assessment; Renoult et al., 2015).

To estimate the perception of *G. cancriformis* color morphs by distinct predators and prey groups, we used the color vision model proposed by Chittka (1992). Although this model has been only validated with behavioral experiments on bees, its general form allow us to apply it for other taxa (e.g. Thery and Casas 2002). There are other models of color vision (Vorobyev and Osorio, 1998; Endler and Mielke, 2005), but when applied correctly, their results tend to be highly correlated (Gawryzewski, 2017). The Chittka (1992) model requires four inputs: (1) the irradiance reaching the observed object, (2) the observer photoreceptor excitation curves, (3) the background reflectance to which photoreceptors are adapted to, and (4) the reflectance curve of the observed object. First, the sensitivity factor *R* was determined for each photoreceptor, as follows:

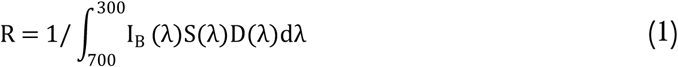

where *I*_B_(λ) is the spectral reflectance function of the background, *S*(λ)is the spectral sensitivity function of each photoreceptor, and *D*(λ) is the illuminant irradiance spectrum. Secondly, the quantum flux *P* (relative amount of photon catch) is calculated, as follows:

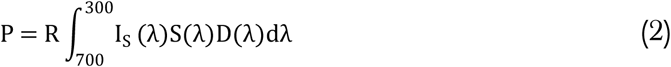

where *I*_S_(λ)is the spectral reflectance function of the stimulus. Assuming that the maximum excitation of a photoreceptor is 1, the phototransduction process is determined by:

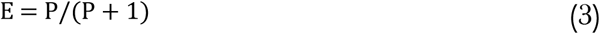

Stimuli spectra are projected in specific color spaces. The coordinates of each spectrum are calculated using photoreceptor excitations, as follows (Chittka et al. 1994):

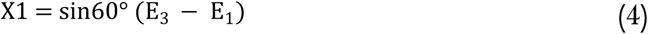

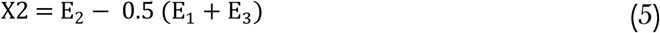

For tetrachromat organisms (Théry and Casas, 2002):

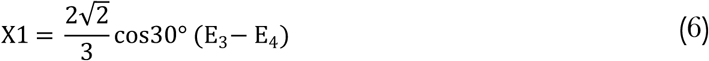

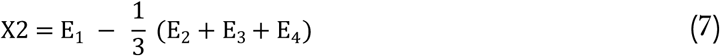

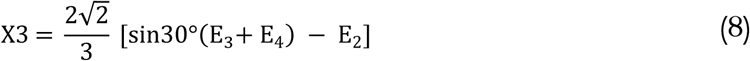

We extended the model of Chittka (1992) to accommodate pentachromatic organisms, as follows:

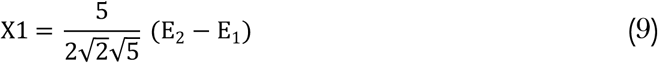

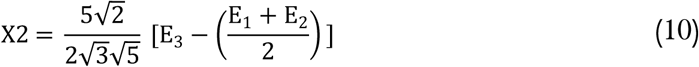

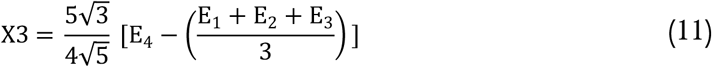

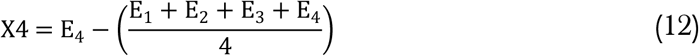

Chromatic contrast between a color stimulus and background, or between two color stimuli, is calculated as the Euclidean distance (Δ*S*) between two points in color space, as follows:

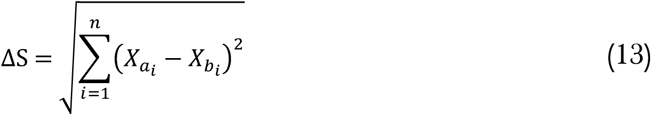

where *X*i (i = 1, 2, 3,…, n) represents the coordinate in the color space.

The color spaces are chromaticity diagrams and, thus, do not estimate the achromatic contrast between objects. Nonetheless, achromatic contrasts can be important visual cues used by both prey and predators. In bees, achromatic contrast is more important than chromatic cues for objects that subtend a visual angle smaller than ∼15°, which means that bees have to be very close to flowers in order to use their color vision for discrimination tasks (Giurfa et al., 1997). Similarly, birds use achromatic contrast in detection of small objects (Osorio et al., 1999). We estimated the achromatic contrast as the excitation (Eq. 3) of the photoreceptor responsible for achromatic discrimination in each organism (Chittka and Kevan 2005).

For our modeling, we used the reflectance data of *G. cancriformis* color morphs that was collected during a previous study (for reflectance curves see figure 1.8 in Gawryszewski, 2007, and figure 5 in Gawryszewski and Motta 2012). These data have already been used to estimate the visual contrast of the yellow, white and the black and white morphs from the perspective of *Apis mellifera* (Gawryszewski and Motta 2012). The spiders were collected from a Brazilian savanna physiognomy, namely Cerrado *sensu stricto*, which is characterized by shrubs and trees of 3 to 8 m tall that are contorted and possess thick, fire-resistant bark, a crown cover of >30%, and additional herbaceous vegetation (Oliveira-Filho and Ratter 2002). The background reflectance was estimated from the average reflectance of leafs, leaf litter, bark, and grasses that were collected from the same area as the spiders (see figure 5 in Gawryszewski and Motta, 2012). To avoid issues with negative values and unrealistic positive values we adjusted the reflectance data by subtracting the reflectance values by the minimum value of each measurement. As illuminant spectrum, we used the International Commission on Illumination (CIE) standard illuminant of D65, which is comparable to open areas, such as the Brazilian savanna (Chittka, 1996).

**Fig. 1.**
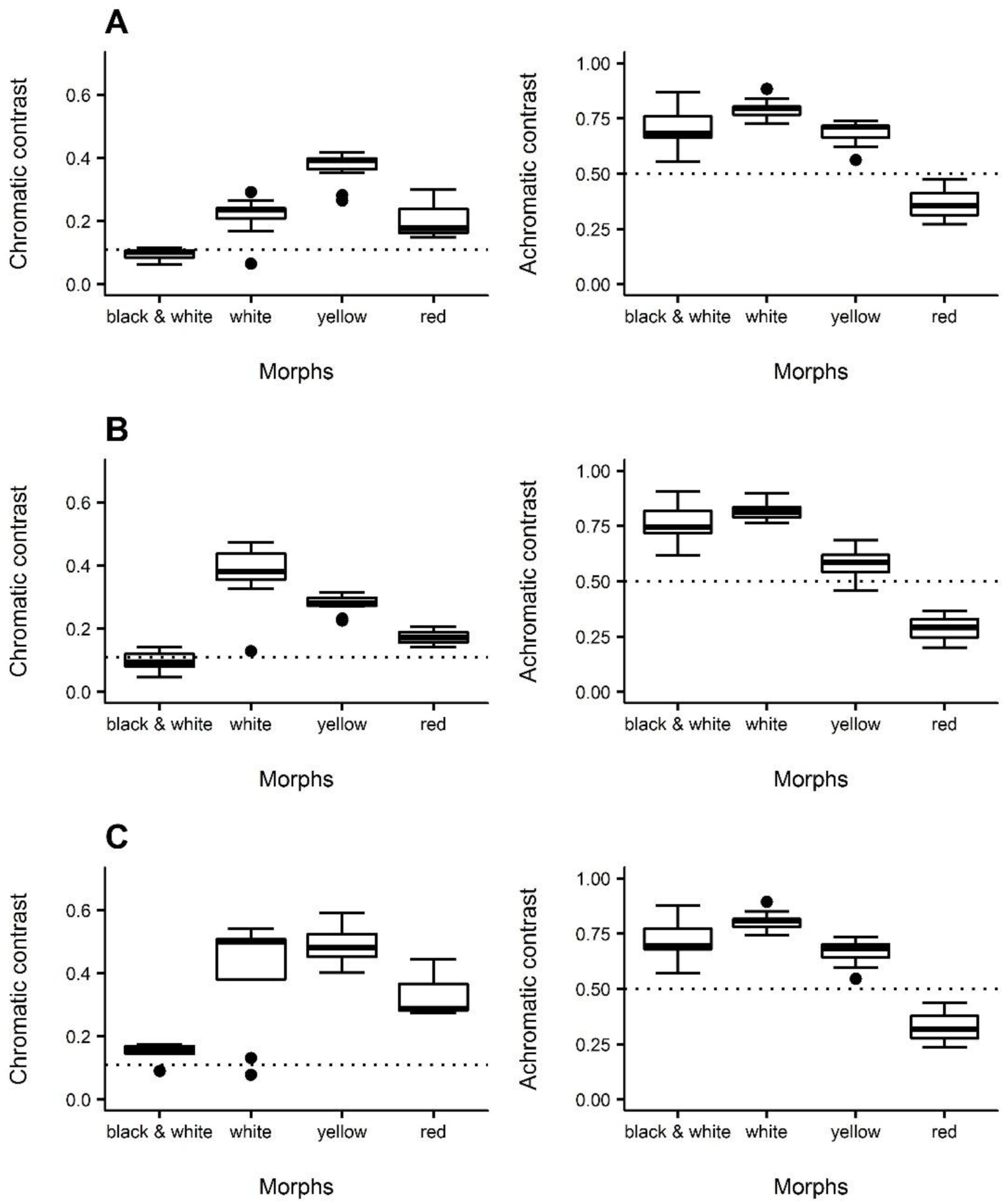
Chromatic (left) and achromatic (right) contrasts of four *Gasteracantha cancriformis* morphs (black and white, N=6; white, N=10; yellow, N=13; and red, N=3) when viewed against a Brazilian savanna background by prey with distinct visual systems. (A) *Apis mellifera* (Hymenoptera). (B) *Drosophila melanogaster* (Diptera). (C) *Fabriciana adippe* (Lepidoptera). Dotted vertical lines represent the discrimination thresholds for chromatic contrast (0.11) and photoreceptor excitation for background in achromatic contrast (0.5).

Visual modeling calculations were conducted using the ‘colourvision’ R package (Gawryszewski, 2017). Linear mixed models were performed using the ‘nlme’ (Pinheiro et al., 2016,) and ‘lme4’ packages (Bates et al. 2015), graphs were plotted using the ‘ggplot2’, ‘ggExtra’, ‘gridExtra’, and ‘pavo’ packages (Wickham, 2009; Maia et al., 2013; Attali, 2016; Auguie, 2016; R Core Team, 2015), and R^2^ values were computed using the package ‘piecewiseSEM’ (Nakagawa and Schielzeth, 2013).

### Multiple prey hypothesis

Using the model described above, we estimated the chromatic and achromatic conspicuousness of the *G. cancriformis* morphs (yellow, white, red, and white patches of the black and white morph) to a suit of potential prey: *A. mellifera* (Hymenoptera, Apidae), *Drosophila melanogaster* (Diptera, Drosophilidae), and *Fabriciana adippe* (Lepidoptera, Nymphalidae). Those species are not necessarily sympatric with *G. cancriformis*. However, these insect orders are commonly intercepted by orb-webs in field experiments (Craig and Ebert 1994; Tso et al. 2002) and represent the diversity of visual systems among insects (Briscoe and Chittka, 2001). Variation within of wavelength of maximum sensitivity is small in Hymenoptera is very little, except for ants and in Lepidoptera most species present four photoreceptor spectral curves (Briscoe and Chittka, 2001). For Diptera, the number of photoreceptors is not so conservative among species and there are not many studies on the color vision of this taxon (Briscoe and Chittka, 2001; Lunau, 2014). Visual modeling work have usually considered Diptera as a tetrachromatic organism (White and Kemp, 2016; White et al., 2016; O’Hanlon et al. 2014). However, recent work showed that *Drosophila melanogaster* use a fifth photoreceptor for color vision (Schnaitmann et al., 2013). Although it remains to be tested whether this species behave as pentachromat we decided to explore this possibility and modelled this is species using five spectral curves for color vision.

For *A. mellifera* and *D. melanogaster*, we used photoreceptor sensitivity curves from the literature (Peitsch et al., 1992; Schnaitmann et al., 2013). It was recently shown that, together with R7-R8 photoreceptors, R1-R6 photoreceptors contribute to color vision in *D. melanogaster* (Kelber and Henze, 2013; Schnaitmann et al., 2013). Therefore, we included the R1-R6 photoreceptor curve, treating *D. melanogaster* as a pentachromat. The graphical curves were extracted directly from the figures of relevant publications using DataThief III version 1.7 (Tummers, 2006). For *Fabriciana adippe*, however, no photoreceptor sensitivity curves are available, so electrophysiological measurements of photoreceptor sensitivity peaks (λ_max;_ Eguchi et al., 1982) were used to estimate the photoreceptor curves (for details see Govardovskii et al., 2000).

For achromatic contrast, bees only use the green photoreceptor (Giurfa et al., 1996), whereas flies only use the outer photoreceptors (R1-R6; Kelber & Henze, 2013). Because the exact mechanism used by lepidopterans for achromatic discrimination is incompletely understood, we assumed that they employ the same mechanism as in bees. The color hexagon model assumes that photoreceptors respond to half their maximum for the background they are adapted to, so that the photoreceptor excitation for background is equivalent to 0.5 units (Chittka, 1992).

The multiple prey hypothesis predicts that different prey taxa perceive color morphs differently. To assess whether each spider morph was perceived differently by prey species, we constructed two linear mixed models, one for chromatic contrast and one for achromatic contrast. Either chromatic (ΔS) or achromatic contrast were used as the dependent variable, and spider morph and prey taxon were used as the independent variables (contrast = spider morph × observer). The spider morph was defined as yellow, white, red, or black and white, and the observers were defined as hymenopteran, dipteran, or lepidopteran. Individual spiders were used as random effects. Normality and homogeneity were verified by visual inspection of quantile-quantile and residuals vs. fitted values plots. We computed all nested models and used the Akaike Information Criterion (AIC) to select the best model. Marginal and conditional R^2^ were estimated according to the recommendations of Nakagawa and Schielzeth (2013).

As a reference point, we used a color discrimination threshold of ΔS= 0.11, which is the threshold value below which trained bees are unable to distinguish different flower colors (Chittka, 1996). However, discrimination thresholds are variable and can change depending on the study species, learning conditions, previous experience, background coloration, whether the task involves discrimination between colors or detection against the background, and whether objects are compared sequentially or simultaneously (Avarguès-Weber & Giurfa, 2014). It should also be noted that threshold values were not behaviorally validated for other taxa.

### Flower mimicry hypothesis

To test the multiple mimic models hypothesis, we compared how flowers and spider morphs are perceived by prey. We gathered all flower reflectance data available in the Floral Reflectance Database (FReD; Arnold et al., 2010), excluding reflectance data from lower flower parts, leaves, bracts, stamens, the inner parts of bell-shaped flowers, and unknown items, as well as spectrum files that did not cover 300 to 700 nm. Most species in the database have only one reflectance spectrum, and for species with multiple reflectance spectra, we randomly selected a single spectrum. We did not average the reflectance of these species because there was no information available on whether these measurements referred to different individuals or different parts of single flowers. In total, we gathered reflectance data from 859 plant species. We grouped flowers visually according to the 10 categories proposed by Chittka et al. (1994), considering whether they reflect or absorb in four spectral ranges, UV (300-400 nm), blue (400-500 nm), green (500-600 nm) and red (600-700 nm). We deleted three spectral curves that did not seem to fit in any of these categories. A caveat of this analysis is that these flowers are not necessarily sympatric to *Gasteracantha cancriformis*. However, flowers spectral curves variation are subtle, because there is a constraint on flower pigments blending (Chittka and Menzel, 1992; Chittka et al. 1994). In addition, we computed reflectance curves from different countries available in FReD database. A qualitative analysis strongly suggests that they all have similar shapes (Fig. S1).

The multiple mimic model hypothesis predicts that different colour morphs are mimicking different flower colors. First, to evaluate color regardless of the observer, we compared hue (Eq. 13), saturation (Eq. 14) and brightness (Eq. 15) of flowers and spiders (Anderson and Prager, 2006):

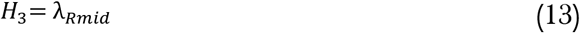

where λ_*Rmid*_ is the wavelength at the middle point between the minimum and maximum reflectances;

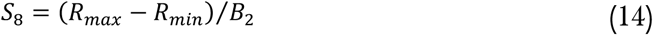

where R_max_ and R_min_ are the maximum and minimum reflectance points; and 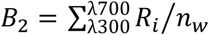, where R_i_ is the reflectance corresponding to each wavelength point, and n_w_ is the total wavelength intervals;

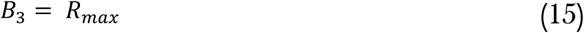

where R_max_ is the maximum reflectance.

We estimated the chromatic difference between individual flowers and the mean achromatic value for each color morph, and calculated the percentage of values below or equal to the theoretical detection threshold of 0.11. Secondly, we computed chromatic distances for spider morphs and flowers following the same steps as for the previous section, considering the visual system of *A. mellifera, D. melanogaster* and *F. adippe*. Then, we calculated a matrix of chromaticity distances between each individual spider color morph and each individual flowers species.

To evaluate if each spider morph and flower category had similar perceptions to each prey species, we constructed two linear mixed models, one for chromatic contrast and one for achromatic contrast. Chromatic or achromatic contrast were used as the dependent variable, and spider morph, flower category and prey taxon were used as the independent variables (contrast = spider morph × flower category × observer). The spider morph was defined as yellow, white, red, or black and white, flower category as ‘1’ to ‘10’, and the observers were defined as hymenopteran, dipteran, or lepidopteran. Individual spiders and individual flowers were considered as random effects. Normality and homogeneity were verified as for the first hypothesis. We selected the best model using AIC, and computed marginal and conditional R^2^ for each model (Nakagawa and Schielzeth, 2013).

As reference points, we used discrimination thresholds of ΔS = 0.11, for the chromatic contrast, and for the achromatic contrast, we assumed the excitation value of 0 for all the three insect taxa.

### Multiple predator hypothesis

The methodology used to investigate the multiple predator hypothesis methodology was very similar to that used for the multiple prey hypothesis, except that we used predator species in our models. As predators, we considered the bird *Parus caeruleus* (Paridae) and the wasp *Philanthus triangulum* (Sphecidae), since birds and wasps are the main predators of orb-web spiders (Rayor, 1996; Foelix, 2010), are visually guided hunters, and have distinct color vision systems. For *P. caeruleus*, we used photoreceptor sensitivity curves available in the literature (Hart, 2001), and for *P. triangulum*, we used photoreceptor sensitivity peaks to estimate photoreceptor sensitivity curves (data available in Briscoe and Chittka, 2001; see Govardovskii et al. 2000 for estimation of sensitivity curves from sensitivity peaks). Again, those species are not sympatric with *G. cancriformis*, but we do not expect a great variation of photoreceptors types within hymenopterans (Peitsch et al., 1992) nor Passeriformes (Hart, 2001).

The multiple predator hypothesis predicts that different predator taxa perceive color morphs differently. To assess this prediction, we established two linear mixed models, one for chromatic contrast and one for achromatic contrast. Either chromatic (ΔS) or achromatic contrast were used as the dependent variable, and spider morph and predator taxon were used as the independent variables (contrast = spider morph × observer). The spider morph was defined as yellow, white, red, or black and white, and individual spiders were used as random effects. Normality and homogeneity were verified by visual inspection of quantile-quantile and residuals vs. fitted values plots. We computed all nested models and used the Akaike Information Criterion to select the best model. We estimated marginal and conditional R^2^ for the models as recommendations of Nakagawa and Schielzeth (2013).

As in the multiple prey hypothesis, we used discrimination thresholds as reference points. For the chromatic contrast, we considered color discrimination thresholds of ΔS = 0.11 and ΔS = 0.06 for the wasp (Dyer and Chittka, 2004) and bird (Théry et al., 2005), respectively. For the achromatic contrast, we considered double cones in birds (Hart, 2001), and assumed green photoreceptors for wasps, as in bees, and compared values obtained to the excitation of 0.5.

## RESULTS

### Multiple prey hypothesis

For chromatic contrast, the model that included the interaction between spider morph and prey taxon presented the lowest AIC value (Table 1). The yellow morph presented the highest ΔS value for *A. mellifera* and *F. adippe* vision, whereas the white spider presented the highest ΔS value for *D. melanogaster*, followed by the yellow morph (Fig. 1). The white patch of the black and white spiders presented a ΔS value that was very close to the theoretical discrimination threshold for all prey species (Fig. 1). The red spiders presented ΔS values near the theoretical discrimination threshold for *A. mellifera* and *D. melanogaster*, but not for *F. adippe* (Fig. 1). For prey achromatic contrast, the model that included the interaction between variables presented the lowest AIC value (Table 1). For all prey groups, the white morph had the highest excitation value, followed by the black and white, yellow, and red morphs, respectively (Fig. 1). The model coefficients are provided in the supplementary material (Table S1 and S2).

**Table 1.** Delta Akaike Information Criterion (ΔAIC) and determination coefficients of Linear Mixed Models of the chromatic and achromatic contrasts of prey and predators.

| Model | df | $\Delta AIC$ | Marginal $R^2$ | Conditional $R^2$ |
| --- | --- | --- | --- | --- |
| ***Multiple prey hypothesis*** |  |  |  |  |
| **Chromatic dimension** |  |  |  |  |
| $\Delta S \sim \text{morph} * \text{observer}$ | 17 | 0.0 | 0.87 | 0.96 |
| $\Delta S \sim \text{morph} + \text{observer}$ | 11 | 23.9 | 0.89 | 0.95 |
| $\Delta S \sim \text{observer}$ | 8 | 52.4 | 0.33 | 0.96 |
| $\Delta S \sim \text{morph}$ | 9 | 61.5 | 0.50 | 0.51 |
| $\Delta S \sim 1$ | 6 | 90.6 | 0 | 0.53 |
| **Achromatic dimension** |  |  |  |  |
| **excitation** $\sim \text{morph} * \text{observer}$ | 17 | 0.0 | 0.82 | 0.99 |
| excitation $\sim \text{morph} + \text{observer}$ | 11 | 57.6 | 0.77 | 0.86 |
| excitation $\sim \text{morph}$ | 9 | 72.2 | 0.79 | 0.89 |
| excitation $\sim \text{observer}$ | 8 | 84.7 | 0.004 | 0.60 |
| excitation $\sim 1$ | 6 | 100.2 | 0 | 0.66 |
| ***Multiple predator hypothesis*** |  |  |  |  |
| **Chromatic dimension** |  |  |  |  |
| $\Delta S \sim \text{morph} * \text{observer}$ | 13 | 0.0 | 0.86 | 0.99 |
| $\Delta S \sim \text{morph} + \text{observer}$ | 10 | 6.9 | 0.88 | 0.99 |
| $\Delta S \sim \text{observer}$ | 7 | 30.6 | 0.29 | 0.99 |
| $\Delta S \sim \text{morph}$ | 9 | 54.5 | 0.58 | 0.58 |
| $\Delta S \sim 1$ | 6 | 74.9 | 0 | 0.63 |
| **Achromatic dimension** |  |  |  |  |
| **excitation ~ morph\*observer** | **14** | **0.0** | **0.78** | **1** |
| excitation ~ morph+observer | 10 | 14.4 | 0.78 | 0.96 |
| excitation ~ observer | 7 | 21.1 | 0.007 | 0.86 |
| excitation ~ morph | 9 | 36.9 | 0.80 | 0.98 |
| excitation ~ 1 | 6 | 43.9 | 0 | 0.92 |

### Flower mimicry hypothesis

We found three peaks of hue for the flowers, around 400, 500 and 600 nm, which are similar to the average hue of spider morphs (Fig. 2A). The saturation metric had only one peak for flowers, to which black and white, white and yellow spider morphs were close (Fig. 2B). The brightness of flowers also only presented a single peak, and white, red and yellow spider morphs had average brightness near to this peak (Fig. 2C).

**Fig. 2.**
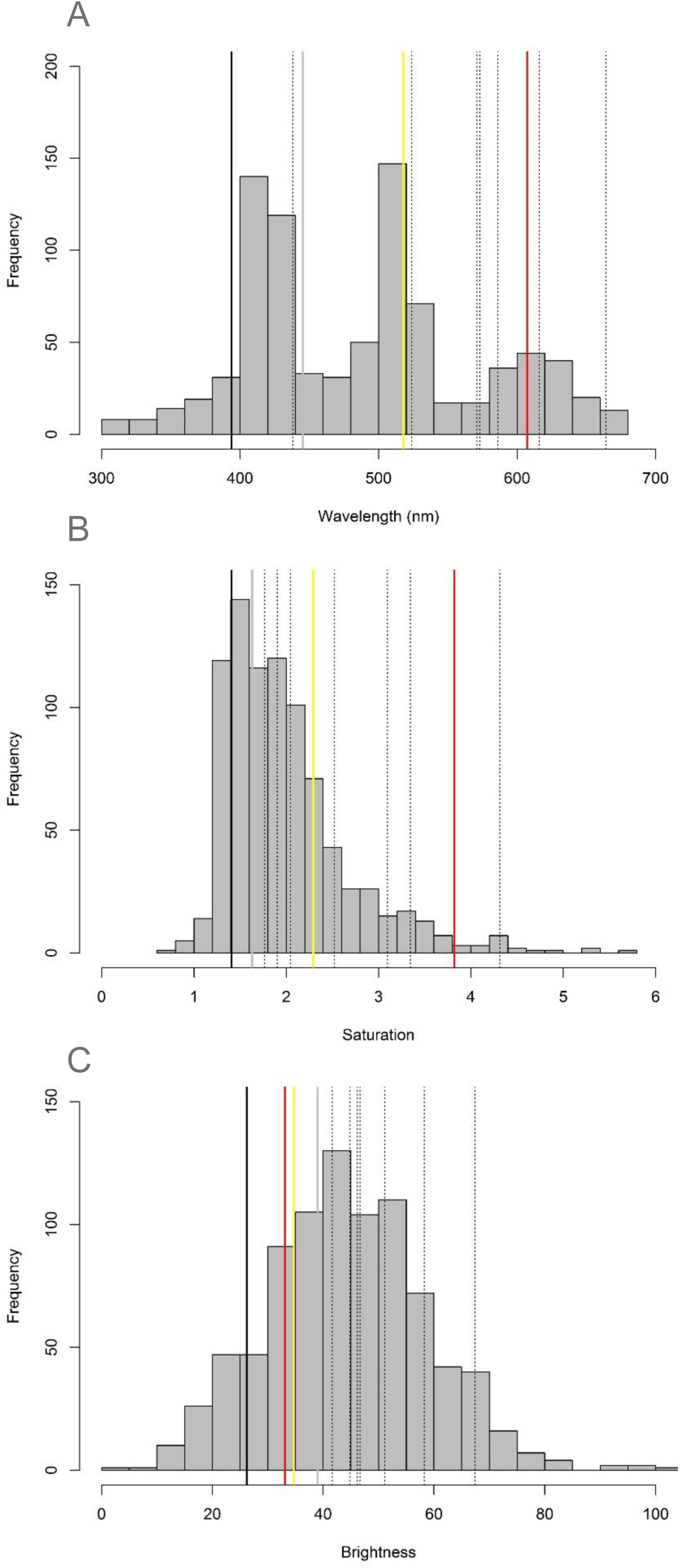
Frequency of color properties of flowers (N = 859): (A) hue, (B) saturation, (C) brightness. Average values of each *Gasteracantha cancriformis* morph are represented with solid colored lines: black and white morph (black line; N = 6), white morph (gray line, N = 10), yellow morph (yellow line, N = 13), red morph (red line, N = 3). Flowers from the Brazilian Savanna (N = 7) are represented with dotted lines.

For all three prey species, only the white patch of the black and white morph had high percentage of values near the chromatic theoretical discrimination threshold of 0.11 when compared to all flowers reflectance spectra: 44.5% for *A. mellifera*, 16.8% for *D. melanogaster*, and 35% for *F. addipe*. For the other spider morphs only a small proportion of the Euclidean distances between flowers and morphs presented values < 0.11. For *A. mellifera* only 1.6% of yellow morphs presented values lower than 0.11, 3.4% of white morphs, and 4.8% of red morphs. For *D. melanogaster* only 2.4% of yellow morphs had values lower than 0.11, 4.0 % of white spiders, and 3.0% of red morphs. For *F. addipe* this values were 0.4%, 0.2%, and 0.5% respectively.

In the color vision model chromatic dimension, the statistical model with interaction among the three variables (flower categories, spider morphs, and prey taxon) had the lowest AIC (Table 2 and S3). In a general view, there seems to be a tendency of growing contrast starting with the black and white morph followed by white, yellow and red morphs (Fig. 3). Only the comparison of black and white spiders and the category ‘8’ of flowers (white flowers that reflect UV) was below the discrimination threshold of 0.11 for all prey taxa (Fig. 3). The categories ‘3’ and ‘4’ compared to black and white spiders were slightly above 0.11 for *A. mellifera*, and ‘4’ for *F. adippe* (Fig. 3). Some categories were around 0.15, which may indicate that for these, flowers and spiders coloration may be perceived as similar to flowers: categories ‘7’ and ‘9’ compared to the black and white morph, and ‘3’ compared to yellow spiders for *A. mellifera*; ‘4’ and ‘9’ compared to the black and white morph for *D. melanogaster*; and ‘3’ compared to black and white, for *F. adippe* (Fig. 3). For the achromatic dimension, the statistical model with interaction among all variables also had the lowest AIC (Table 2 and S4). Most of the groups had excitation values around 0 and 0.2, regardless the spider morph and observer (Fig. 3).

**Table 2.** Delta Akaike Information Criterion (ΔAIC) and determination coefficients of Linear Mixed Models of the chromatic and achromatic contrasts between spider morphs and flower categories.

| | df | $\Delta AIC$ | Marg.<br>R<sup>2</sup> | Cond.<br>R<sup>2</sup> |
| --- | --- | --- | --- | --- |
| **Chromatic dimension** |  |  |  |  |
| $\Delta S \sim \text{morph} * \text{flower} * \text{observer} + (1 \vert \text{IDflower}) + (1 \vert \text{IDspider})$ | 111 | 0 | 0.67 | 0.82 |
| $\Delta S \sim \text{morph} * \text{flower} + \text{observer} + (1 \vert \text{IDflower}) + (1 \vert \text{IDspider})$ | 41 | -22276.39 | 0.60 | 0.75 |
| $\Delta S \sim \text{morph} * \text{flower} + (1 \vert \text{IDflower}) + (1 \vert \text{IDspider})$ | 39 | -39316.11 | 0.55 | 0.70 |
| $\Delta S \sim \text{morph} * \text{observer} + \text{flower} + (1 \vert \text{IDflower}) + (1 \vert \text{IDspider})$ | 23 | -58911.74 | 0.46 | 0.61 |
| $\Delta S \sim \text{morph} * \text{observer} + (1 \vert \text{IDflower}) + (1 \vert \text{IDspider})$ | 15 | -59819.05 | 0.28 | 0.61 |
| $\Delta S \sim \text{morph} + \text{flower} * \text{observer} + (1 \vert \text{IDflower}) + (1 \vert \text{IDspider})$ | 33 | -62659.22 | 0.44 | 0.59 |
| $\Delta S \sim \text{Flower} * \text{observer} + (1 \vert \text{IDflower}) + (1 \vert \text{IDspider})$ | 30 | -62679.69 | 0.25 | 0.60 |
| $\Delta S \sim \text{morph} + \text{flower} + \text{observer} + (1 \vert \text{IDflower}) + (1 \vert \text{IDspider})$ | 17 | -64825.9 | 0.43 | 0.58 |
| $\Delta S \sim \text{flower} + \text{observer} + (1 \vert \text{IDflower}) + (1 \vert \text{IDspider})$ | 14 | -64846.37 | 0.24 | 0.58 |
| $\Delta S \sim \text{morph} + \text{observer} + (1 \vert \text{IDflower}) + (1 \vert \text{IDspider})$ | 9 | -65733.21 | 0.25 | 0.58 |
| $\Delta S \sim \text{observer} + (1 \vert \text{IDflower}) + (1 \vert \text{IDspider})$ | 6 | -65753.68 | 0.06 | 0.57 |
| $\Delta S \sim \text{morph} + \text{flower} + (1 \vert \text{IDflower}) + (1 \vert \text{IDspider})$ | 15 | -75210.75 | 0.37 | 0.52 |
| $\Delta S \sim \text{flower} + (1 \vert \text{IDflower}) + (1 \vert \text{IDspider})$ | 12 | -75231.22 | 0.18 | 0.52 |
| $\Delta S \sim \text{morph} + (1 \vert \text{IDflower}) + (1 \vert \text{IDspider})$ | 7 | -76118.06 | 0.20 | 0.52 |
| $\Delta S \sim 1 + (1 \vert \text{IDflower}) + (1 \vert \text{IDspider})$ | 4 | -76138.53 | 0 | 0.52 |
| **Achromatic dimension** |  |  |  |  |
| excitation ~ morph * flower * observer + (1 \| IDflower) + (1 \| IDspider) | 111 | 0 | 0.49 | 0.64 |
| excitation ~ morph * flower + observer + (1 \| IDflower) + (1 \| IDspider) | 41 | -13215.1 | 0.42 | 0.57 |
| excitation ~ morph * flower + (1 \| IDflower) + (1 \| IDspider) | 39 | -14202.51 | 0.42 | 0.57 |
| excitation ~ morph + flower * observer + (1 \| IDflower) + (1 \| IDspider) | 33 | -33305.63 | 0.30 | 0.45 |
| excitation ~ morph * observer + flower + (1 \| IDflower) + (1 \| IDspider) | 23 | -33320.44 | 0.30 | 0.44 |
| excitation ~ Flower * observer + (1 \| IDflower) + (1 \| IDspider) | 30 | -33331.64 | 0.12 | 0.45 |
| excitation ~ morph * observer + (1 \| IDflower) + (1 \| IDspider) | 15 | -33835.46 | 0.20 | 0.45 |
| excitation ~ morph + flower + observer + (1 \| IDflower) + (1 \| IDspider) | 17 | -35420.86 | 0.29 | 0.43 |
| excitation ~ flower + observer + (1 \| IDflower) + (1 \| IDspider) | 14 | -35446.87 | 0.11 | 0.43 |
| excitation ~ morph + observer + (1 \| IDflower) + (1 \| IDspider) | 9 | -35935.88 | 0.18 | 0.43 |
| excitation ~ observer + (1 \| IDflower) + (1 \| IDspider) | 6 | -35961.89 | 0.05 | 0.43 |
| excitation ~ morph + flower + (1 \| IDflower) + (1 \| IDspider) | 15 | -36160.68 | 0.28 | 0.43 |
| excitation ~ flower + (1 \| IDflower) + (1 \| IDspider) | 12 | -36186.69 | 0.10 | 0.43 |
| excitation ~ morph + (1 \| IDflower) + (1 \| IDspider) | 7 | -36675.7 | 0.18 | 0.43 |
| excitation ~ 1 + (1 \| IDflower) + (1 \| IDspider) | 4 | -36701.71 | 0 | 0.43 |

**Fig. 3.**
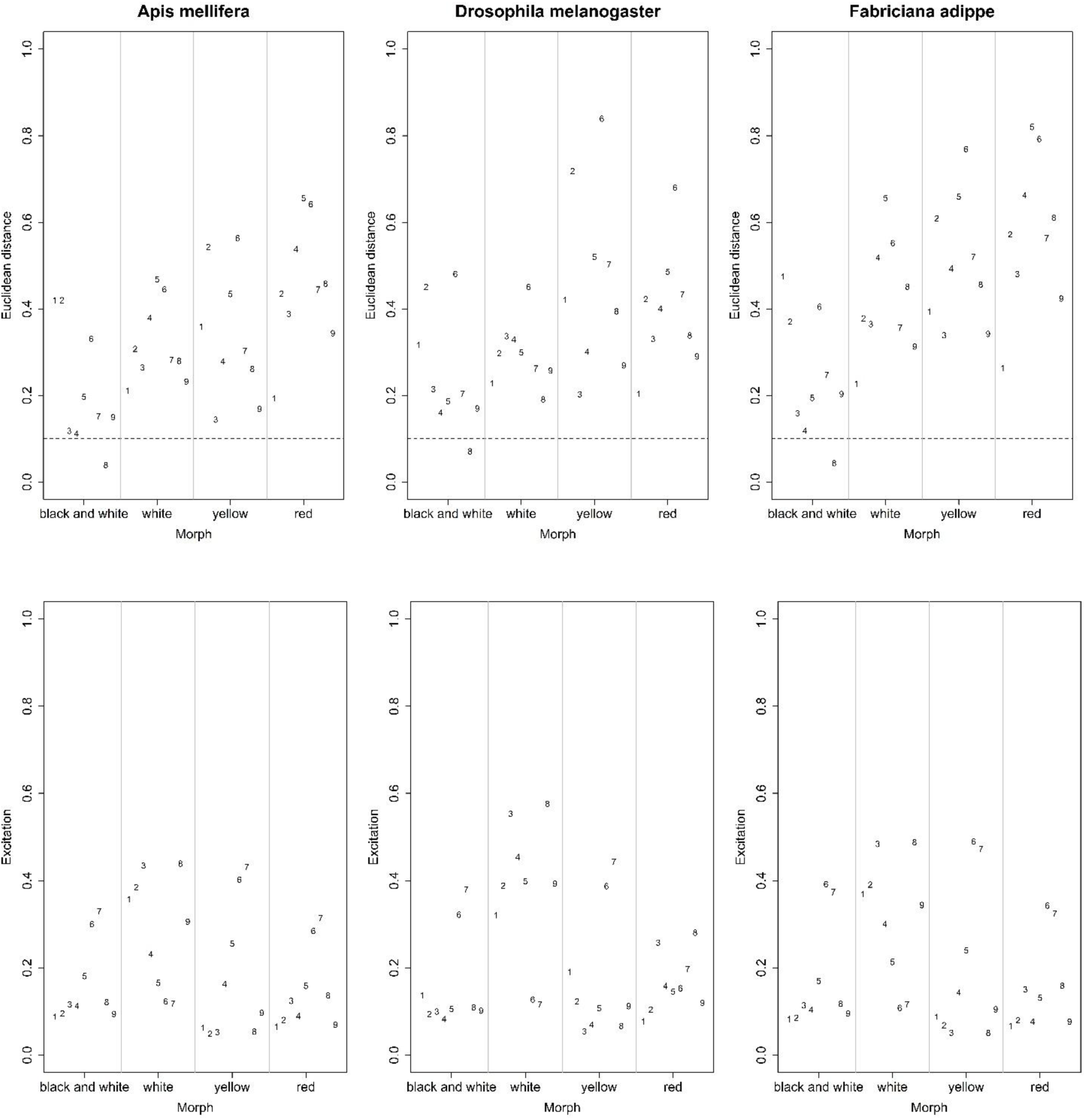
Chromatic (upper) and achromatic (lower) contrasts of four *Gasteracantha cancriformis* morphs (black and white, N=6; white, N=10; yellow, N=13; and red, N=3) when compared with ten flowers categories (Chittka et al., 1994) indicated by numbers. These values were computed considering the Brazilian savanna as background and based on three potencial prey: *Apis mellifera* (Hymenoptera; left), *Drosophila melanogaster* (Diptera; middle), and *Fabriciana adippe* (Lepidoptera; right). Dotted vertical lines represent the discrimination thresholds for chromatic contrast (0.11).

### Multiple predator hypothesis

For the chromatic contrast, the model with interaction between variables presented the lowest AIC value (Table 1). The black and white morph presented the lowest ΔS value for both predators (Fig. 4A,B; Table S5). The white morph was the one with highest ΔS value for *P. caeruleus*, though yellow and red morphs presented similar values (Fig. 4A). For *P. triangulum*, the white spider morph presented the highest ΔS value, followed by the yellow and red morphs. The latter was near the theoretical discrimination threshold of 0.11 (Fig. 4B). For the achromatic contrast, the model that included the interaction between variables presented the lowest AIC value (Table 1), even though the values of the two predator species were very similar. For *P. caeruleus*, the white morph had the highest excitation value, followed by the yellow, black and white, and red morphs, respectively (Fig. 4C). The white morph also had the highest excitation value for *P. triangulum*, followed by the black and white, yellow, and red morphs, respectively (Fig. 4D). The model coefficients are provided in the supplementary material (Table S5 and S6).

**Fig. 4.**
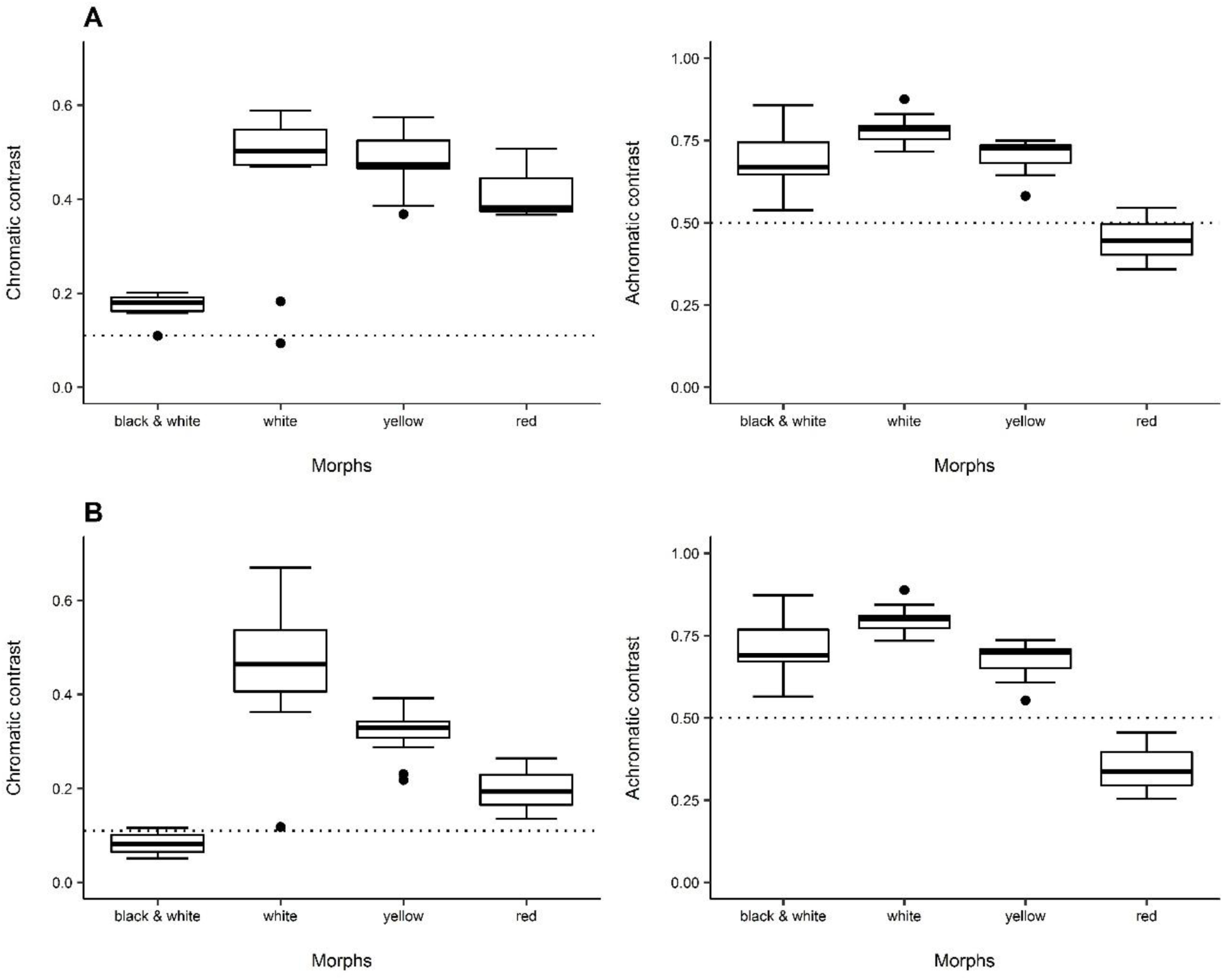
Chromatic (left) and achromatic (right) contrasts of four *Gasteracantha cancriformis* morphs (black and white, N=6; white, N=10; yellow, N=13; and red, N=3) when viewed against a Brazilian savanna background by predators with distinct visual systems. (A) *Parus caeruleus* (Passeriformes). (B) *Philanthus Triangulum* (Hymenoptera). Dotted vertical lines represent the discrimination thresholds for chromatic contrast (0.06) and photoreceptor excitation for background in achromatic contrast (0.5).

## DISCUSSION

Our statistical analyses show that the majority of *G. cancriformis* morphs have a high probability of being detected by potential prey and the degree of detectability varies according to the receiver. Some spider morphs are also conspicuous for predators, but the multiple predator hypothesis was partially corroborated, because the degree of detectability between predators was similar. In addition, we offer some support for the flower mimicry hypothesis.

### Multiple prey hypothesis

In *G. cancriformis*, spider morphs conspicuousness is perceived differently by prey species. The yellow and white morphs were the most conspicuous to all prey taxa. The former being more contrasting from the background for honeybee color vision, and the latter, for flies. The red morph, although inconspicuous for honeybee and flies, showed high detectability for butterflies. A recent study using the receptor noise-limited color vision model showed that insects prey perceive coloration of *Verrucosa arenata* morphs differently, however the maintenance of color polymorphism does not seem to be influenced by multiple prey as we suggested here. Yellow morphs of *V. arenata* have higher chromatic contrast than white morphs for Diptera and Hymenoptera. Whereas in the achromatic dimension the white morph had a higher contrast for both prey taxa (Ajuria-Ibarra et al. 2017). Color morphs may, instead be influenced by other factors such as different visual channels of relevant observers or illumination (Ajuria-Ibarra et al. 2017; White and Kemp, 2016).

The hypothesis that different morphs exploit different visual channels in prey was also proposed in another study to explain the evolution of color polymorphism in *Gasteracantha fornicata*. The yellow morphs of *G. fornicata* would benefit from stimulating the dipteran chromatic channel, whereas white morphs would benefit from stimulating the achromatic channel (White and Kemp, 2016). For the achromatic dimension, although the statistical analyses also suggested an interaction between spider morph and prey taxon, spider morphs presented similar levels of achromatic detectability comparing among prey taxa, therefore, this idea seems inconsistent with the multiple prey hypothesis for *G. cancriformis.* However, when comparing chromatic and achromatic contrasts of each prey taxa individually, we observe different detectabilities between the two visual channels for the morphs. Therefore, the hypothesis of exploitation of different visual channels of prey could be possible to explain the color polymorphism, as in White & Kemp (2016), but possibly not enough to explain such a diverse color variation as occur in *G. cancriformis*.

### Flower mimicry hypothesis

Several authors have proposed flower mimicry hypothesis as a mechanism of prey attraction by orb-web spiders *via* conspicuous body coloration (e.g. Craig and Ebert, 1994; Hauber, 2002). However, the hypothesis has seldom been tested. Flower mimicry using color vision modelling has been tested for orchid mantis (*Hymenopus coronatus*) prey (O’Hanlon et al., 2013). Color vision modelling suggested that pollinators are unable to distinguish the colors of the mantis and flowers, and a field experiment showed that the mantis actually attracts more pollinators than flowers (O’Hanlon et al., 2013). Our results showed that, considering only color metrics, most of the *G. cancriformis* morphs indeed have similar coloration to flowers. However, when we modeled color perception to potential prey, only the black and white morphs is similar to a category of white flowers. Similarly to our results, in *G. fornicata* the white morphs seems to be indistinguishable from sympatric flowers according to results of bee color vision modeling, but yellow morphs and flowers were not perceptually different (Maia & White, 2017). Conversely, a study of various orb-web spider species that also used color vision models found that, as perceived by dipterans and hymenopterans, the colors of spiders are very similar to those of flowers (White et al., 2016). Both pieces of evidence are circumstantial. They may only reflect the diversity of flower colors and spider colors. Additionally, in the Brazilian savanna, *G. cancriformis* is abundant during the transition from the wet to the dry season, which overlaps partially the flowering period of woody plants (Oliveira, 1998; Gouveia and Fefili, 1998). Flowering peaks in these plants is related to pollinators occurrence, that is around April to October (Oliveira, 1998; Gouveia and Fefili, 1998). Therefore, flower coloration mimicry would be an advantageous foraging strategy to spiders that are abundant during this period of the year. However, a field experiment conducted with *G. cancriformis* showed that color had no effect on prey capture success (Gawryzewsky and Motta, 2012). Furthermore, it could be possible that insects do not represent a strong selection force, considering that most of taxa only perceive chromatic contrast when they are very close to the object (Giurfa et al., 1997).

### Multiple predators hypothesis

The results of the present study do not strongly support the multiple predator hypothesis for the maintenance of color polymorphism in *G. cancriformis*, as the spider morphs present the same order of conspicuousness in both the chromatic and achromatic dimensions. Even so, red morphs are particularly more conspicuous to a bird than to a wasp. Therefore, this signal could be targeting bird predators but would appear relatively inconspicuous to a hymenopteran predator and prey. In contrast, the white and yellow morphs are highly detectable by both predator taxa. The colors of two of the four *G. cancriformis* morphs (yellow and red) are typical of aposematic species (Endler and Mappes, 2004). Conspicuous coloration is especially advantageous when it increases the mismatch with the background and facilitates predator learning (Endler and Greenwood, 1988). Spiders of the genus *Gasteracantha* possess spines and a hard abdomen. Moreover, the hunting wasp *Sceliphron laetum* avoids provisioning initial instars with *Gasteracantha* spiders (Elgar and Jebb, 1999). Morphological and behavioral defenses that make ingestion difficult along with the species’ bright colors constitute aposematism (Endler and Greenwood, 1988; Ruxton et al., 2004). Though aposematism is not commonly reported in spiders (Oxford and Gillespie, 1998), Brandley et al. (2016) conducted an experiment with black widow models and found that models with red markings were more likely to be avoided by birds than all black models.

Color polymorphism may seem counterintuitive in aposematic species, but it may occur when frequency-dependent selection is different among predators, for instance, when a predator presents strong apostatic selection, while other predator has a strong anti-apostatic selection (Endler and Greenwood, 1988). It is also possible when there is a covariance between the relative crypsis of morphs for one predator and frequency-dependent selection. However, contrarily to the first scenario, the equilibrium is unstable (Endler and Greenwood, 1988). Besides, scenarios of overdominance, or equal fitness from different selection pressures may also influence (Stevens and Ruxton, 2012). In *G. cancriformis*, morphs have variable degrees of conspicuousness for a single predator or for multiple predators, therefore they might be subject to distinct types of selection.

Not only selective pressures from prey and predators may influence color polymorphism, but also thermoregulatory effects and the effect of illumination on the signaler detectability (Rao and Mendoza-Cuenca, 2016; Rojas et al., 2014). Therefore, polymorphism may result from multiple evolutionary forces, in which some morphs signals their impalatability to predators, whereas other morphs are protected from certain predators due to camouflage, meanwhile, they may benefit from thermoregulatory behavior by occupying different microhabitats.

Most of studies focus on a single signal receiver, however, we could better understand signal evolution if we considered that individuals interact with different kinds of observers, whether they are mutualists or antagonists (Endler and Mappes, 2014; Schaefer et al., 2004). The multiple receiver hypothesis has been evaluated in intersexual and intrasexual relations (Guindre-Parker et al., 2012), signaler interaction with prey and predators (Endler, 1983), and interaction with pollinators and herbivores (Irwin et al., 2003). For instance, in the snow bunting (*Plectrophenax nivalis*), multiple achromatic patches signal distinct information to females and males: wing coloration inform about male immune response and reproductive performance, whereas plumage of the rectrices and mantle convey information about territoriality and probable aggression (Guindre-Parker et al., 2012). Multiple receivers also maintain guppy color polymorphism, males can have black, orange, yellow or iridescent spots, that influence on female attraction, but they vary in frequency and size accordingly with predation risk (Endler, 1983). Lastly, flowers polymorphism is also influenced by multiple receivers as in the wild radish (*Raphanus sativus*). Plants that produce anthocyanin – a defense component - and plants that do not produce vary in coloration. Therefore, herbivores may use coloration as cue to find the anthocyanin-recessive morphs (Irwin et al., 2003).

Here, we present a small step of the multiple receivers hypothesis on the evolution of color polymorphism, multiple functions may also maintain this variation, although it remains to be tested. Variation of signal receivers alone may not be sufficient to explain color polymorphism, and gene flow may act together on the maintenance of color variation (Gray and McKinnon, 2007). We only considered chromatic and achromatic discrimination, but color pattern geometry, shape, contour, size, angle, texture, and distance of visual detection (Troscianko et al., 2009) may also influence the behavior of both prey and predators toward spiders since different species use distinct visual cues for stimuli detection and recognition (Théry and Gomez, 2010). Furthermore, color vision models do not include other perceptual mechanisms, such as cognition, color categorization, past experiences, or memory imprecision (Renoult et al., 2015), even though these factors may affect detectability and, consequently, influence the survival rate of morphs in different ways (Théry and Gomez, 2010). Additionally, non-adaptive explanations, such as overdominance and allele equilibrium in absence of selection, are often ignored when studying polymorphisms in an ecological perspective. Finally, predation experiments, field experiments that evaluate prey taxa caught by the different spider morphs, and ecological data on abudance and composition of prey and predators populations that occur sympatrically with *G. cancriformis* are paramount to validate and complement the findings of the present study.

## Acknowledgments

We thank CAPES for financial support (CAPES/PROEX), and for a scholarship awarded to NXG. Prof. Rodrigo Willemart, Prof. Fausto Nomura, and anonymous reviewers for their comments on the manuscript.

## Author contribution

NXG and FMG contributed to the design of the study. NXG wrote the manuscript and ran the statistical analyses. FMG supervised the analyses and commented on the manuscript.

