## Supplementary Materials for "Orb-web spider color polymorphism through the eyes of multiple prey and predators"

### Supplementary information

### Tables

Table S1. Results of Linear Mixed Model comparing chromatic contrast of *Gasteracantha cancriformis* morphs viewed by different groups of prey.

| | Value | SE | df | t | p |
|---|---|---|---|---|---|
| Intercept | 0.0947 | 0.0175 | 56 | 5.4185 | < 0.0001 |
| Red morph | 0.1136 | 0.0436 | 28 | 2.6019 | 0.0146 |
| White morph | 0.1221 | 0.0365 | 28 | 3.3455 | 0.0023 |
| Yellow morph | 0.2773 | 0.0221 | 28 | 12.5034 | < 0.0001 |
| Observer *F. adippe* | 0.0540 | 0.0072 | 56 | 7.4013 | < 0.0001 |
| Observer *D. melanogaster* | 0.0020 | 0.0072 | 56 | 0.2712 | 0.7872 |
| Red morph & observer *F. adippe* | 0.0736 | 0.0462 | 56 | 1.5913 | 0.1172 |
| White morph & observer *F. adippe* | 0.1334 | 0.0421 | 56 | 3.1700 | 0.0024 |
| Yellow morph & observer *F. adippe* | 0.0662 | 0.0130 | 56 | 5.1003 | < 0.0001 |
| Red morph & observer *D. melanogaster* | -0.0375 | 0.0462 | 56 | -0.8121 | 0.4202 |
| White morph & observer *D. melanogaster* | 0.1530 | 0.0421 | 56 | 3.6366 | 0.0006 |
| Yellow morph & observer *D. melanogaster* | -0.0936 | 0.0130 | 56 | -7.2091 | < 0.0001 |

Table S2. Results of Linear Mixed Model (log transformed) comparing achromatic contrast of *Gasteracantha cancriformis* morphs viewed by different groups of prey.

| | Value | SE | df | t | p |
|---|---|---|---|---|---|
| Intercept | -0.3605 | 0.0491 | 56 | -7.3419 | <0.0001 |
| Red morph | -0.67 | 0.0869 | 28 | -7.7143 | <0.0001 |
| White morph | 0.1291 | 0.0620 | 28 | 2.0817 | 0.0466 |
| Yellow morph | -0.0224 | 0.0598 | 28 | -0.3742 | 0.7110 |
| Observer *F. adippe* | 0.0799 | 0.0074 | 56 | 10.7462 | <0.0001 |
| Observer *D. melanogaster* | 0.0205 | 0.0074 | 56 | 2.7551 | 0.0079 |
| Red morph & observer *F. adippe* | -0.3309 | 0.0281 | 56 | -11.7844 | <0.0001 |
| White morph & observer *F. adippe* | -0.0513 | 0.0080 | 56 | -6.4438 | <0.0001 |
| Yellow morph & observer *F. adippe* | -0.2531 | 0.0133 | 56 | -19.0999 | <0.0001 |
| Red morph & observer *D. melanogaster* | -0.1307 | 0.0281 | 56 | -4.6552 | <0.0001 |
| White morph & observer *D. melanogaster* | -0.0042 | 0.0080 | 56 | -0.5228 | 0.6032 |
| Yellow morph & observer *D. melanogaster* | -0.0466 | 0.0133 | 56 | -3.5162 | 0.0009 |

Table S3. Results of Linear Mixed Model comparing chromatic contrast between *Gasteracantha cancriformis* morphs and flowers categories viewed by different prey taxa.

| Model | Estimate | SE | t |
|---|---|---|---|
| Intercept | 0.6484 | 0.0197 | 32.8746 |
| Red morph | -0.1878 | 0.0330 | -5.6836 |
| White morph | -0.0489 | 0.0241 | -2.0245 |
| Yellow morph | -0.2082 | 0.0231 | -9.0274 |
| Flower A2 | 0.0001 | 0.0083 | 0.0150 |
| Flower A3 | -0.3043 | 0.0069 | -43.8400 |

| | | | |
|---|---|---|---|
| Flower A4 | -0.3132 | 0.0072 | -43.4090 |
| Flower A5 | -0.2044 | 0.0081 | -25.3850 |
| Flower A6 | -0.0728 | 0.0143 | -5.08680 |
| Flower A7 | -0.2588 | 0.0114 | -22.7270 |
| Flower A8 | -0.4495 | 0.0174 | -25.7980 |
| Flower A9 | -0.2612 | 0.0098 | -26.5480 |
| Observer Drosophila melanogaster | -0.0850 | 0.0044 | -19.2410 |
| Observer Fabriciana adippe | 0.0408 | 0.0044 | 9.2319 |
| Red morph & Flower A2 | 0.0942 | 0.0076 | 12.4097 |
| White morph & Flower A2 | 0.1376 | 0.0055 | 24.8206 |
| Yellow morph & Flower A2 | 0.2194 | 0.0053 | 41.4260 |
| Red morph & Flower A3 | 0.3584 | 0.0064 | 56.2490 |
| White morph & Flower A3 | 0.0850 | 0.0047 | 18.2793 |
| Yellow morph & Flower A3 | 0.4878 | 0.0044 | 109.685 |
| Red morph & Flower A4 | 0.4685 | 0.0066 | 70.7186 |
| White morph & Flower A4 | 0.2424 | 0.0048 | 50.1102 |
| Yellow morph & Flower A4 | 0.6069 | 0.0046 | 131.2700 |
| Red morph & Flower A5 | 0.4284 | 0.0074 | 57.9372 |
| White morph & Flower A5 | 0.2641 | 0.0054 | 48.9139 |
| Yellow morph & Flower A5 | 0.5742 | 0.0052 | 111.245 |
| Red morph & Flower A6 | 0.2795 | 0.0131 | 21.2845 |
| White morph & Flower A6 | 0.2238 | 0.0096 | 23.3318 |
| Yellow morph & Flower A6 | 0.4338 | 0.0092 | 47.3292 |
| Red morph & Flower A7 | 0.3302 | 0.0105 | 31.5845 |
| White morph & Flower A7 | 0.2096 | 0.0076 | 27.4618 |
| Yellow morph & Flower A7 | 0.4857 | 0.0073 | 66.5636 |
| Red morph & Flower A8 | 0.5181 | 0.0160 | 32.3841 |
| White morph & Flower A8 | 0.3614 | 0.0117 | 30.9283 |
| Yellow morph & Flower A8 | 0.6866 | 0.0112 | 61.4827 |
| Red morph & Flower A9 | 0.2827 | 0.0090 | 31.3022 |
| White morph & Flower A9 | 0.0729 | 0.0066 | 11.0498 |
| Yellow morph & Flower A9 | 0.4074 | 0.0063 | 64.6294 |
| Red morph & Observer Drosophila melanogaster | 0.1025 | 0.0076 | 13.3999 |
| White morph & Observer Drosophila melanogaster | 0.1351 | 0.0056 | 24.1782 |
| Yellow morph & Observer Drosophila melanogaster | 0.0962 | 0.0053 | 18.0101 |
| Red morph & Observer Fabriciana adippe | -0.0247 | 0.0076 | -3.2256 |
| White morph & Observer Fabriciana adippe | -0.0123 | 0.0056 | -2.2106 |
| Yellow morph & Observer Fabriciana adippe | 0.0326 | 0.0053 | 6.1073 |
| Flower A2 & Observer Drosophila melanogaster | 0.1079 | 0.0062 | 17.4204 |
| Flower A3 & Observer Drosophila melanogaster | 0.2044 | 0.0052 | 39.2915 |
| Flower A4 & Observer Drosophila melanogaster | 0.1504 | 0.0054 | 27.8129 |
| Flower A5 & Observer Drosophila melanogaster | 0.0733 | 0.0060 | 12.1394 |
| Flower A6 & Observer Drosophila melanogaster | 0.2028 | 0.0107 | 18.9168 |
| Flower A7 & Observer Drosophila melanogaster | 0.1475 | 0.0085 | 17.2773 |

| | | | |
|---|---|---|---|
| Flower A8 & Observer Drosophila melanogaster | 0.1530 | 0.0131 | 11.7110 |
| Flower A9 & Observer Drosophila melanogaster | 0.1105 | 0.0074 | 14.9859 |
| Flower A2 & Observer Fabriciana adippe | -0.0804 | 0.0062 | -12.9670 |
| Flower A3 & Observer Fabriciana adippe | 0.0141 | 0.0052 | 2.7043 |
| Flower A4 & Observer Fabriciana adippe | -0.0317 | 0.0054 | -5.8550 |
| Flower A5 & Observer Fabriciana adippe | -0.0436 | 0.0060 | -7.2254 |
| Flower A6 & Observer Fabriciana adippe | 0.0200 | 0.0107 | 1.8699 |
| Flower A7 & Observer Fabriciana adippe | 0.0674 | 0.0085 | 7.8984 |
| Flower A8 & Observer Fabriciana adippe | -0.0306 | 0.0131 | -2.3404 |
| Flower A9 & Observer Fabriciana adippe | 0.0232 | 0.0074 | 3.1434 |
| Red morph & Flower A2 & Observer Drosophila melanogaster | -0.1347 | 0.0107 | -12.553 |
| White morph & Flower A2 & Observer Drosophila melanogaster | -0.0476 | 0.0078 | -6.0668 |
| Yellow morph & Flower A2 & Observer Drosophila melanogaster | -0.1288 | 0.0075 | -17.1900 |
| Red morph & Flower A3 & Observer Drosophila melanogaster | -0.1558 | 0.0090 | -17.2910 |
| White morph & Flower A3 & Observer Drosophila melanogaster | -0.1843 | 0.0066 | -28.0100 |
| Yellow morph & Flower A3 & Observer Drosophila melanogaster | -0.2642 | 0.0063 | -42.0040 |
| Red morph & Flower A4 & Observer Drosophila melanogaster | -0.2096 | 0.0094 | -22.3780 |
| White morph & Flower A4 & Observer Drosophila melanogaster | -0.1804 | 0.0068 | -26.3750 |
| Yellow morph & Flower A4 & Observer Drosophila melanogaster | -0.2624 | 0.0065 | -40.1270 |
| Red morph & Flower A5 & Observer Drosophila melanogaster | -0.2279 | 0.0105 | -21.7880 |
| White morph & Flower A5 & Observer Drosophila melanogaster | -0.0612 | 0.0076 | -8.0132 |
| Yellow morph & Flower A5 & Observer Drosophila melanogaster | -0.1972 | 0.0073 | -27.0170 |
| Red morph & Flower A6 & Observer Drosophila melanogaster | -0.2162 | 0.0186 | -11.642 |
| White morph & Flower A6 & Observer Drosophila melanogaster | -0.0871 | 0.0136 | -6.4241 |
| Yellow morph & Flower A6 & Observer Drosophila melanogaster | -0.1904 | 0.0130 | -14.6900 |
| Red morph & Flower A7 & Observer Drosophila melanogaster | -0.1846 | 0.0148 | -12.487 |
| White morph & Flower A7 & Observer Drosophila melanogaster | -0.039 | 0.0108 | -3.6120 |
| Yellow morph & Flower A7 & Observer Drosophila melanogaster | -0.1675 | 0.0103 | -16.2280 |
| Red morph & Flower A8 & Observer Drosophila melanogaster | -0.2625 | 0.0226 | -11.6000 |
| White morph & Flower A8 & Observer Drosophila melanogaster | -0.0862 | 0.0165 | -5.2161 |
| Yellow morph & Flower A8 & Observer Drosophila melanogaster | -0.2592 | 0.0158 | -16.413 |
| Red morph & Flower A9 & Observer Drosophila melanogaster | -0.1026 | 0.0128 | -8.0349 |
| White morph & Flower A9 & Observer Drosophila melanogaster | -0.0525 | 0.0093 | -5.6302 |
| Yellow morph & Flower A9 & Observer Drosophila melanogaster | -0.1692 | 0.0089 | -18.9770 |
| Red morph & Flower A2 & Observer Fabriciana adippe | 0.1247 | 0.0107 | 11.6173 |
| White morph & Flower A2 & Observer Fabriciana adippe | 0.0955 | 0.0078 | 12.1804 |
| Yellow morph & Flower A2 & Observer Fabriciana adippe | 0.1036 | 0.0075 | 13.8307 |
| Red morph & Flower A3 & Observer Fabriciana adippe | 0.0594 | 0.0090 | 6.5938 |
| White morph & Flower A3 & Observer Fabriciana adippe | 0.1597 | 0.0066 | 24.2682 |
| Yellow morph & Flower A3 & Observer Fabriciana adippe | -0.0176 | 0.0063 | -2.8060 |
| Red morph & Flower A4 & Observer Fabriciana adippe | 0.1204 | 0.0094 | 12.8481 |
| White morph & Flower A4 & Observer Fabriciana adippe | 0.1771 | 0.0068 | 25.8879 |
| Yellow morph & Flower A4 & Observer Fabriciana adippe | 0.0386 | 0.0065 | 5.9057 |
| Red morph & Flower A5 & Observer Fabriciana adippe | 0.1531 | 0.0105 | 14.6420 |

| | | | |
|---|---|---|---|
| White morph & Flower A5 & Observer Fabriciana adippe | 0.1680 | 0.0076 | 22.0036 |
| Yellow morph & Flower A5 & Observer Fabriciana adippe | 0.0661 | 0.0073 | 9.0582 |
| Red morph & Flower A6 & Observer Fabriciana adippe | 0.0398 | 0.0186 | 2.1433 |
| White morph & Flower A6 & Observer Fabriciana adippe | 0.0781 | 0.0136 | 5.7571 |
| Yellow morph & Flower A6 & Observer Fabriciana adippe | -0.0044 | 0.0130 | -0.3422 |
| Red morph & Flower A7 & Observer Fabriciana adippe | -0.0185 | 0.0148 | -1.2520 |
| White morph & Flower A7 & Observer Fabriciana adippe | 0.0749 | 0.0108 | 6.9422 |
| Yellow morph & Flower A7 & Observer Fabriciana adippe | -0.0574 | 0.0103 | -5.5642 |
| Red morph & Flower A8 & Observer Fabriciana adippe | 0.1573 | 0.0226 | 6.9528 |
| White morph & Flower A8 & Observer Fabriciana adippe | 0.1665 | 0.0165 | 10.0765 |
| Yellow morph & Flower A8 & Observer Fabriciana adippe | 0.0617 | 0.0158 | 3.9048 |
| Red morph & Flower A9 & Observer Fabriciana adippe | 0.0381 | 0.0128 | 2.9857 |
| White morph & Flower A9 & Observer Fabriciana adippe | 0.1223 | 0.0093 | 13.1130 |
| Yellow morph & Flower A9 & Observer Fabriciana adippe | -0.0317 | 0.0089 | -3.5558 |

Table S4. Results of Linear Mixed Model comparing achromatic contrast between *Gasteracantha cancriformis* morphs and flowers categories viewed by different prey taxa.

| Model | Estimate | SE | t |
|---|---|---|---|
| Intercept | 0.2971 | 0.0173 | 17.2212 |
| Red morph | 0.3008 | 0.0282 | 10.6557 |
| White morph | -0.0468 | 0.0206 | -2.2721 |
| Yellow morph | -0.0412 | 0.0197 | -2.0915 |
| Flower A2 | 0.0126 | 0.0101 | 1.2565 |
| Flower A3 | 0.0441 | 0.0084 | 5.2143 |
| Flower A4 | 0.0397 | 0.0088 | 4.5255 |
| Flower A5 | 0.1285 | 0.0098 | 13.1099 |
| Flower A6 | 0.2509 | 0.0174 | 14.4121 |
| Flower A7 | 0.2771 | 0.0139 | 19.9995 |
| Flower A8 | 0.0516 | 0.0212 | 2.4312 |
| Flower A9 | 0.0105 | 0.0120 | 0.8747 |
| Observer Drosophila melanogaster | 0.0733 | 0.0062 | 11.7617 |
| Observer Fabriciana adippe | -0.0093 | 0.0062 | -1.4998 |
| Red morph & Flower A2 | 0.0100 | 0.0107 | 0.9353 |
| White morph & Flower A2 | -0.0415 | 0.0078 | -5.3009 |
| Yellow morph & Flower A2 | 0.0161 | 0.0075 | 2.1574 |
| Red morph & Flower A3 | 0.0172 | 0.0090 | 1.9142 |
| White morph & Flower A3 | -0.0642 | 0.0066 | -9.7760 |
| Yellow morph & Flower A3 | 0.0528 | 0.0063 | 8.4106 |
| Red morph & Flower A4 | -0.1563 | 0.0093 | -16.7157 |
| White morph & Flower A4 | 0.1142 | 0.0068 | 16.7301 |
| Yellow morph & Flower A4 | 0.0041 | 0.0065 | 0.6263 |
| Red morph & Flower A5 | -0.3195 | 0.0104 | -30.6121 |
| White morph & Flower A5 | 0.1266 | 0.0076 | 16.6108 |
| Yellow morph & Flower A5 | 0.0141 | 0.0073 | 1.9307 |

| | | | |
|---|---|---|---|
| Red morph & Flower A6 | -0.4980 | 0.0185 | -26.8691 |
| White morph & Flower A6 | 0.1334 | 0.0135 | 9.8533 |
| Yellow morph & Flower A6 | 0.0269 | 0.0129 | 2.0781 |
| Red morph & Flower A7 | -0.5311 | 0.0148 | -35.9991 |
| White morph & Flower A7 | 0.1295 | 0.0108 | 12.0209 |
| Yellow morph & Flower A7 | 0.0276 | 0.0103 | 2.6848 |
| Red morph & Flower A8 | 0.0132 | 0.0226 | 0.5862 |
| White morph & Flower A8 | -0.0696 | 0.0165 | -4.2205 |
| Yellow morph & Flower A8 | 0.0627 | 0.0158 | 3.9759 |
| Red morph & Flower A9 | -0.0550 | 0.0127 | -4.3123 |
| White morph & Flower A9 | 0.0509 | 0.0093 | 5.4676 |
| Yellow morph & Flower A9 | -0.0019 | 0.0089 | -0.2127 |
| Red morph & Observer Drosophila melanogaster | -0.1048 | 0.0108 | -9.7119 |
| White morph & Observer Drosophila melanogaster | 0.1136 | 0.0079 | 14.4037 |
| Yellow morph & Observer Drosophila melanogaster | -0.0502 | 0.0075 | -6.6681 |
| Red morph & Observer Fabriciana adippe | 0.0197 | 0.0108 | 1.8248 |
| White morph & Observer Fabriciana adippe | 0.0564 | 0.0079 | 7.1593 |
| Yellow morph & Observer Fabriciana adippe | 0.0124 | 0.0075 | 1.6509 |
| Flower A2 & Observer Drosophila melanogaster | -0.0761 | 0.0087 | -8.7013 |
| Flower A3 & Observer Drosophila melanogaster | -0.0982 | 0.0073 | -13.3800 |
| Flower A4 & Observer Drosophila melanogaster | -0.1219 | 0.0076 | -15.9634 |
| Flower A5 & Observer Drosophila melanogaster | -0.1735 | 0.0085 | -20.3588 |
| Flower A6 & Observer Drosophila melanogaster | -0.0536 | 0.0151 | -3.5409 |
| Flower A7 & Observer Drosophila melanogaster | -0.0311 | 0.0120 | -2.5776 |
| Flower A8 & Observer Drosophila melanogaster | -0.0911 | 0.0184 | -4.9412 |
| Flower A9 & Observer Drosophila melanogaster | -0.0612 | 0.0104 | -5.8834 |
| Flower A2 & Observer Fabriciana adippe | -0.0070 | 0.0087 | -0.8007 |
| Flower A3 & Observer Fabriciana adippe | 0.0062 | 0.0073 | 0.8500 |
| Flower A4 & Observer Fabriciana adippe | -0.0049 | 0.0076 | -0.6431 |
| Flower A5 & Observer Fabriciana adippe | -0.0036 | 0.0085 | -0.4260 |
| Flower A6 & Observer Fabriciana adippe | 0.0877 | 0.0151 | 5.7935 |
| Flower A7 & Observer Fabriciana adippe | 0.0466 | 0.0120 | 3.8648 |
| Flower A8 & Observer Fabriciana adippe | 0.0039 | 0.0184 | 0.2100 |
| Flower A9 & Observer Fabriciana adippe | 0.0111 | 0.0104 | 1.0701 |
| Red morph & Flower A2 & Observer Drosophila melanogaster | 0.1106 | 0.0151 | 7.2993 |
| White morph & Flower A2 & Observer Drosophila melanogaster | 0.0187 | 0.0111 | 1.6891 |
| Yellow morph & Flower A2 & Observer Drosophila melanogaster | 0.0923 | 0.0106 | 8.7274 |
| Red morph & Flower A3 & Observer Drosophila melanogaster | 0.2144 | 0.0127 | 16.8616 |
| White morph & Flower A3 & Observer Drosophila melanogaster | -0.0846 | 0.0093 | -9.1141 |
| Yellow morph & Flower A3 & Observer Drosophila melanogaster | 0.2304 | 0.0089 | 25.9555 |
| Red morph & Flower A4 & Observer Drosophila melanogaster | 0.3463 | 0.0132 | 26.1874 |
| White morph & Flower A4 & Observer Drosophila melanogaster | -0.2045 | 0.0097 | -21.1812 |
| Yellow morph & Flower A4 & Observer Drosophila melanogaster | 0.1975 | 0.0092 | 21.3991 |
| Red morph & Flower A5 & Observer Drosophila melanogaster | 0.4296 | 0.0148 | 29.1033 |

| | | | |
|---|---|---|---|
| White morph & Flower A5 & Observer Drosophila melanogaster | -0.1909 | 0.0108 | -17.7114 |
| Yellow morph & Flower A5 & Observer Drosophila melanogaster | 0.1337 | 0.0103 | 12.9770 |
| Red morph & Flower A6 & Observer Drosophila melanogaster | 0.0913 | 0.0262 | 3.4847 |
| White morph & Flower A6 & Observer Drosophila melanogaster | -0.1453 | 0.0191 | -7.5887 |
| Yellow morph & Flower A6 & Observer Drosophila melanogaster | -0.1116 | 0.0183 | -6.1022 |
| Red morph & Flower A7 & Observer Drosophila melanogaster | 0.0601 | 0.0209 | 2.8818 |
| White morph & Flower A7 & Observer Drosophila melanogaster | -0.1466 | 0.0152 | -9.6195 |
| Yellow morph & Flower A7 & Observer Drosophila melanogaster | -0.1084 | 0.0146 | -7.4422 |
| Red morph & Flower A8 & Observer Drosophila melanogaster | 0.2192 | 0.0319 | 6.8630 |
| White morph & Flower A8 & Observer Drosophila melanogaster | -0.0692 | 0.0233 | -2.9655 |
| Yellow morph & Flower A8 & Observer Drosophila melanogaster | 0.2280 | 0.0223 | 10.2286 |
| Red morph & Flower A9 & Observer Drosophila melanogaster | 0.1665 | 0.018 | 9.2373 |
| White morph & Flower A9 & Observer Drosophila melanogaster | -0.1020 | 0.0132 | -7.7481 |
| Yellow morph & Flower A9 & Observer Drosophila melanogaster | 0.1201 | 0.0126 | 9.5430 |
| Red morph & Flower A2 & Observer Fabriciana adippe | 0.0015 | 0.0151 | 0.1022 |
| White morph & Flower A2 & Observer Fabriciana adippe | 0.0010 | 0.0111 | 0.0937 |
| Yellow morph & Flower A2 & Observer Fabriciana adippe | 0.0021 | 0.0106 | 0.2015 |
| Red morph & Flower A3 & Observer Fabriciana adippe | 0.0201 | 0.0127 | 1.5824 |
| White morph & Flower A3 & Observer Fabriciana adippe | -0.0565 | 0.0093 | -6.0851 |
| Yellow morph & Flower A3 & Observer Fabriciana adippe | 0.0264 | 0.0089 | 2.9726 |
| Red morph & Flower A4 & Observer Fabriciana adippe | 0.0621 | 0.0132 | 4.6940 |
| White morph & Flower A4 & Observer Fabriciana adippe | -0.0675 | 0.0097 | -6.9939 |
| Yellow morph & Flower A4 & Observer Fabriciana adippe | -0.0196 | 0.0092 | -2.1285 |
| Red morph & Flower A5 & Observer Fabriciana adippe | 0.0485 | 0.0148 | 3.2861 |
| White morph & Flower A5 & Observer Fabriciana adippe | -0.0587 | 0.0108 | -5.4486 |
| Yellow morph & Flower A5 & Observer Fabriciana adippe | -0.0354 | 0.0103 | -3.4344 |
| Red morph & Flower A6 & Observer Fabriciana adippe | -0.1194 | 0.0262 | -4.5546 |
| White morph & Flower A6 & Observer Fabriciana adippe | -0.0694 | 0.0191 | -3.6279 |
| Yellow morph & Flower A6 & Observer Fabriciana adippe | -0.0387 | 0.0183 | -2.1146 |
| Red morph & Flower A7 & Observer Fabriciana adippe | -0.0596 | 0.0209 | -2.8573 |
| White morph & Flower A7 & Observer Fabriciana adippe | -0.0630 | 0.0152 | -4.1340 |
| Yellow morph & Flower A7 & Observer Fabriciana adippe | -0.0408 | 0.0146 | -2.8010 |
| Red morph & Flower A8 & Observer Fabriciana adippe | 0.0218 | 0.0319 | 0.6811 |
| White morph & Flower A8 & Observer Fabriciana adippe | -0.0567 | 0.0233 | -2.4308 |
| Yellow morph & Flower A8 & Observer Fabriciana adippe | 0.0221 | 0.0223 | 0.9916 |
| Red morph & Flower A9 & Observer Fabriciana adippe | 0.0120 | 0.0180 | 0.6637 |
| White morph & Flower A9 & Observer Fabriciana adippe | -0.0455 | 0.0132 | -3.4530 |
| Yellow morph & Flower A9 & Observer Fabriciana adippe | -0.0014 | 0.0126 | -0.1094 |

Table S5. Results of Linear Mixed Model comparing chromatic contrast of *Gasteracantha cancriformis* morphs viewed by different groups of predators.

| | Value | SE | df | t | p |
|---|---|---|---|---|---|
| Intercept | 0.1704 | 0.0220 | 28 | 7.7355 | < 0.0001 |
| Red morph | 0.2484 | 0.0413 | 28 | 6.0153 | < 0.0001 |

| | | | | | |
|---|---|---|---|---|---|
| White morph | 0.2761 | 0.0510 | 28 | 5.4093 | < 0.0001 |
| Yellow morph | 0.3163 | 0.0267 | 28 | 11.8502 | < 0.0001 |
| Observer *P. triangulum* | -0.0873 | 0.0077 | 28 | -11.394 | < 0.0001 |
| Red morph & observer *P. triangulum* | -0.1333 | 0.0260 | 28 | -5.1233 | < 0.0001 |
| White morph & observer *P. triangulum* | 0.0964 | 0.0613 | 28 | 1.5726 | 0.1270 |
| Yellow morph & observer *P. triangulum* | -0.0794 | 0.0096 | 28 | -8.2352 | < 0.0001 |

Table S6. Results of Linear Mixed Model (log transformed) comparing achromatic contrast of *Gasteracantha cancriformis* morphs viewed by different groups of predators.

| | Value | SE | df | t | p |
|---|---|---|---|---|---|
| Intercept | -0.3809 | 0.0428 | 28 | -8.898 | < 0.0001 |
| Red morph | -0.4321 | 0.0741 | 28 | -5.8287 | < 0.0001 |
| White morph | 0.1362 | 0.0541 | 28 | 2.5159 | 0.0179 |
| Yellow morph | 0.0246 | 0.0517 | 28 | 0.4763 | 0.6376 |
| Observer *P. triangulum* | 0.0333 | 0.0047 | 28 | 7.1148 | < 0.0001 |
| Red & observer *P. triangulum* | -0.3015 | 0.0484 | 28 | -6.2265 | < 0.0001 |
| White & observer *P. triangulum* | -0.0124 | 0.0048 | 28 | -2.5765 | 0.0155 |
| Yellow & observer *P. triangulum* | -0.0742 | 0.007 | 28 | -10.601 | < 0.0001 |

### Figures

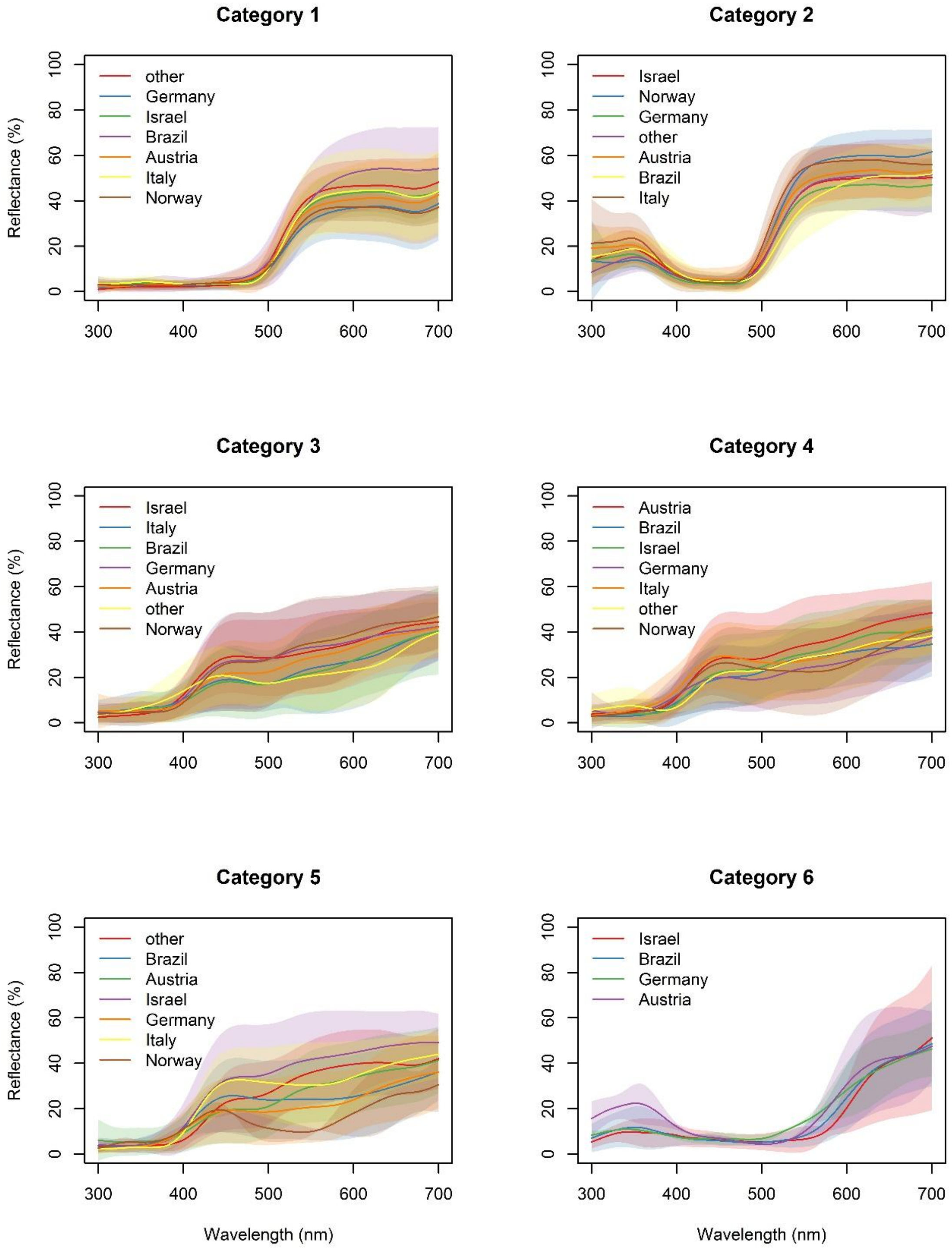

Category 1
other
Germany
Israel
Brazil
Austria
Italy
Norway
Reflectance (%)
0
20
40
60
80
100
300
400
500
600
700
Category 2
Israel
Norway
Germany
other
Austria
Brazil
Italy
Category 3
Israel
Italy
Brazil
Germany
Austria
other
Norway
Category 4
Austria
Brazil
Israel
Germany
Italy
other
Norway
Category 5
other
Brazil
Austria
Israel
Germany
Italy
Norway
Category 6
Israel
Brazil
Germany
Austria
Wavelength (nm)

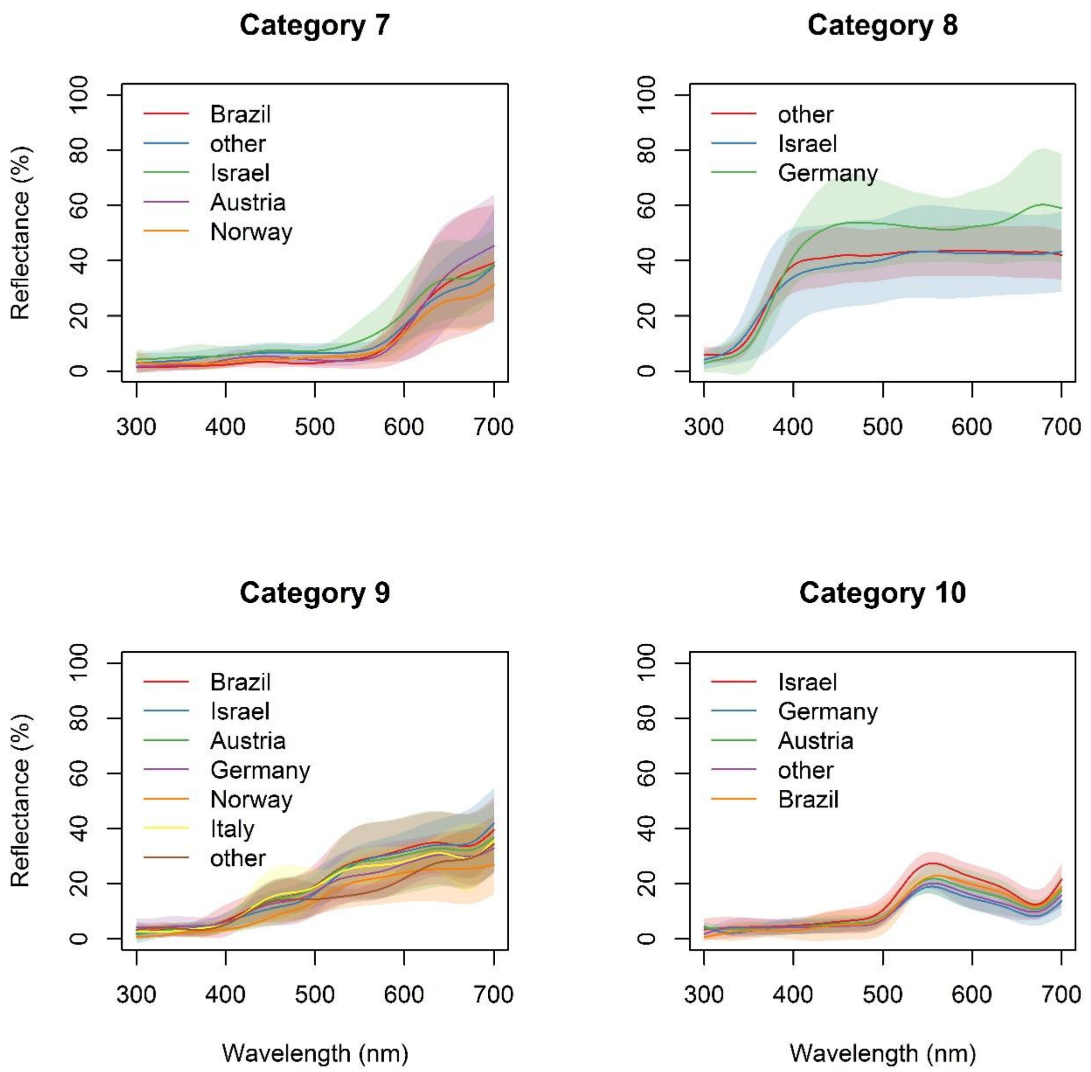


Fig S1. Reflectance spectra of flowers from the Floral Reflectance Database (FReD), grouped according to ten categories proposed by Chittka et al. (1994) and countries where they were colected: category 1, N= 94; category 2, N = 97; category 3, N = 240; category 4, N = 187; category 5, N = 108; category 6, N = 20; category 7, N = 35; category 8, N = 13; category 9, N = 53; category 10, N = 17. Countries with only one reflectance curve per category were grouped into the category ‘other’.
